# The autophagy core protein Atg18 regulates exosome release and rescues pathogenic dysfunction associated to Parkinson Sac domain mutation in Synaptojanin

**DOI:** 10.64898/2026.05.26.727892

**Authors:** Irene Sanchez-Mirasierra, Sergio Hernández-Díaz, Liam Barry-Carroll, Aida Arjona Martí, Saurav Ghimire, Jean-Christophe Delpech, Sandra-Fausia Soukup

**Affiliations:** Univ. Bordeaux, CNRS, IMN, UMR 5293, F-33000 Bordeaux, France; Univ. Bordeaux, INRAE, Bordeaux INP, NutriNeuro, 33000, Bordeaux, France; Multidisciplinary Institute of Ageing, MIA-Portugal, University of Coimbra, 3004-504, Coimbra, Portugal; Achucarro Basque Center for Neuroscience, E-48940 Leioa, Spain; Ikerbasque Basque Foundation for Science, E-48009 Bilbao, Spain

**Keywords:** Exosome release, Autophagy, Synaptojanin, Multivesicular Bodies, Drosophila, Parkinson’s disease, Locomotion, Presynaptic terminals

## Abstract

Macroautophagy, a catabolic process conserved across evolution, participates in fundamental aspects of synaptic function and neuronal survival. Exosomes are extracellular vesicles that mediate cell communication, including emerging roles mediating brain intercellular signaling, synaptic function and neuronal survival. Indeed, both autophagy and exosome release interact with each other, but this interplay is poorly characterized at the neuronal synapse, especially in the context of neurodegenerative diseases. Here, we report that at the presynaptic compartment the autophagy protein Atg18a/WIPI2 regulates exosome release at the level of the multivesicular body. The Parkinson disease mutation R258Q in the protein Synaptojanin inhibits synaptic autophagy, and causes locomotion deficits, seizures and neurodegeneration. We found that this mutation also reduces exosome release and that the dopaminergic overexpression of Atg18a in Synaptojanin mutant animals restores exosome release, locomotion and dopaminergic survival without restoring synaptic autophagy. Our data reports a novel function of Atg18a in the regulation of exosome release and spotlights the role of exosome release in the pathogenesis of Parkinson disease.

## INTRODUCTION

Autophagy is a catabolic pathway essential for neuronal survival and for the degradation and recycling of proteins, lipids and organelles. At the presynaptic compartment, synaptic autophagy functions in the maintenance of synaptic homeostasis by degrading components thereby regulating not only local proteostasis, but also neurotransmission, synapse development and plasticity (Gupta et al., 2016; Hernandez-Diaz et al., 2022; Hernandez et al., 2012; Hoffmann-Conaway et al., 2020; Kiral et al., 2020). Interestingly, autophagy is regulated locally by presynaptic enriched proteins in response to metabolic as well as neuronal stimuli (Bademosi et al., 2023; Soukup et al., 2016; Vanhauwaert, Kuenen, Masius, Bademosi, Manetsberger, Schoovaerts, Bounti, Gontcharenko, Swerts, Vilain, Picillo, Barone, Munshi, Vrij, Kushner, Gounko, Mandemakers, Bonifati, Meunier, Soukup, Verstreken, et al., 2017). To identify how autophagy regulates other processes linked to brain homeostasis and communication seems essential to fully understand the functions of autophagy in brain health and disease. Exosomes are extracellular vesicles, derived from endocytic compartments, that can transport RNA species (e.g. miRNA), DNA, proteins and lipids *via* the extracellular space (Chivet et al., 2013; Miranda et al., 2018; Prada et al., 2018) (Ghanam et al., 2022). Exosomes have been described as a mechanism to mediate cell-cell communication (Chivet et al., 2013). Intriguingly, evidence supports yet not well understood emerging roles of exosomes in neurological diseases (Grad et al., 2014; Papadopoulos et al., 2018; Rajendran et al., 2006; Wang et al., 2017) and as regulators of synaptic activity and plasticity, neurogenesis, nerve regeneration and neuronal survival, function and viability (Chiang et al., 2021; Ching & Kingham, 2015; Frühbeis et al., 2013; Lachenal et al., 2011; Sharma et al., 2019; Spelat et al., 2022). Moreover, exosomes may also function as a waste disposal mechanism in the CNS (Chen et al., 2001; Lopez-Verrilli et al., 2013; Yuyama et al., 2015) highlighting the role of exosomes as a regulators of neuronal health. Alterations of the endolysosomal and autophagy systems are common in neurodegenerative diseases (Aman et al., 2021; Friedman et al., 2012; Hernandez-Diaz & Soukup, 2020; Komatsu et al., 2006; Liang et al., 2010; Nixon & Rubinsztein, 2024; Sanchez-Mirasierra et al., 2022) and these perturbations can impact the content of exosomes by resorting dysfunctional and potential toxic proteins (Izco et al., 2022; Miranda et al., 2018). Synapses are the communication hubs of the neuron. A precise regulation of protein supply together with the elimination of nonfunctional or toxic proteins is necessary to sustain synaptic function during neuronal communication. Intriguingly, synaptic accumulation of toxic proteins and synaptic dysfunction are reported in neurodegenerative diseases preceding neuronal loss (Burke & O’Malley, 2013; Cheng et al., 2010; Hornykiewicz, 1998; Kramer & Schulz-Schaeffer, 2007). These observations support the idea that autophagy and exosome release may cooperate to maintain brain homeostasis and neuronal function and survival. In contrast, autophagy impairment could spoil exosome release (number and/or content) worsening neuronal survival and contributing to the progression of neurodegenerative diseases (Xu et al., 2018). Synaptic autophagy is an emerging field far from being fully understood. In particular, the crosstalk of autophagy with exosome release at the synapse is not well characterized and whether deregulation of this interplay at the synapse plays a role during the onset and progression of neurodegeneration is mostly unknown. Interestingly, the subcellular origin of exosomes could determine the fate and the cargo of the exosomes (Yokoi et al., 2019), but how exosomes of synaptic origin could fulfill functions linked to synaptic function or neuronal survival is not well characterized. Indeed, molecular mechanisms linking exosome release and synaptic autophagy, especially during pathological scenarios are currently missing.

There are cellular components that are common to both autophagosome and exosome biogenesis (Puri et al., 2018; Xu et al., 2018; Zubkova et al., 2024). For instance, the endolysosomal system is central for exosome biogenesis but also participates in autophagy. This is further supported by the observation that the Multivesicular Body (MVB), a specialized form of late endosome where protein sorting takes place, can engage both autophagy and exosome release pathways. MVB actually participates in autophagy during the maturation of the autophagosome (Fader, Sánchez, et al., 2008) while the fusion of the MVB, that contain intraluminal vesicles, with the plasma membrane also results in the release of the intraluminal vesicles as exosomes into the extracellular space. In non-neuronal cells, induction of autophagy using amino acid deprivation seems to inhibit exosome release while inhibition of autophagy favors that process (Fader, Sanchez, et al., 2008; Villarroya-Beltri et al., 2016). However, scientific literature also reports scenarios where inhibition of autophagy in hepatocytes actually blocks exosome release (Shrivastava et al., 2016). Thus, the effect of autophagy on exosome release is still unclear and seems to be very context specific.

Presynaptic terminals depend for their correct functioning on efficient protein sorting and recycling. Furthermore, evidence indicates that synaptic dysfunction precedes neuronal loss in many neurodegenerative diseases. Unfortunately, the interplay of autophagy and exosome release at the synapse during neurodegeneration or cell-cell communication remains still largely enigmatic. Moreover, little is known about how the deregulation of synaptic autophagy affects exosome release and whether dysfunction of the exosome-autophagy interplay could be at the root of neurodegeneration since detailed studies about the regulation of this interplay in neurons and in the presynaptic compartment are still missing.

Here we show that autophagy and exosome release are interconnected at the presynaptic terminal in healthy condition, and identify a novel function of the autophagy core protein Atg18a, the invertebrate homologue of WIPI2, in the biogenesis of multivesicular bodies and exosomes. Moreover, in pathological condition, we show that increasing Atg18a protein levels rescues altered exosome release, locomotion, seizure-like behaviors and dopaminergic neurodegeneration in animals harboring a Parkinson mutation in the synaptic protein Synaptojanin (Synj^R258Q^). This work connects for the first time how autophagic alterations affect exosome release at the synapse and explore the consequences of this interplay in the context of Parkinson’s disease.

## RESULTS

### Autophagy regulates exosome release from presynaptic terminals

The relatively small size of exosomes and the complex anatomy of the nervous system makes the analysis of exosome release particularly challenging. However, the neuronal presynaptic compartment of the neuromuscular junction (NMJ) in *Drosophila* larvae is very well differentiated from the muscular postsynaptic compartment, making the NMJ an excellent model system to study and quantify exosome release from the neuronal presynaptic compartment (Sanchez-Mirasierra et al., 2021). To monitor exosome release, we expressed the exosome marker Evenness Interrupted fused with GFP (Evi::GFP) specifically in motor neuron and marked the presynaptic area with an antibody (anti HRP) (**Fig 1A,A’,** magenta) that recognizes neuronal membranes in *Drosophila* (Jan & Jan, 1982). Using the UAS-Gal4 expression system and the motor neurons specific driver (D42-Gal4) allows specific expression in motor neurons on the presynaptic side with no expression in the muscle on the postsynaptic side. The Evi:GFP signal detected inside of the presynapse correspond to the intraluminal vesicles (precursors of the exosomes) contained within the Multi Vesicular Body (MVB) that, upon the fusion with the cytoplasmic membrane, gives rise to the released exosomes. Accordingly, we can also clearly detect Evi::GFP signal outside of the synaptic membrane labeling the exosomes released into the extracellular space (**Fig 1A”**, yellow line marks limit of HRP signal). To analyze exosome release we quantified Evi::GFP fluorescence in the extracellular space at the NMJ with an automatic quantification method that we previously published (Sanchez-Mirasierra et al., 2021). We first converted the confocal image to a binary image and applied a threshold to mark presynaptic areas (anti-HRP signal) in a single plane. Then we calculated the surrounding area in a radius of 1µm, corresponding to the postsynaptic area where the exosomes are released and created a mask for the exosome release site. This mask was applied on the EVI::GFP image with the exosome marker. As a result, only the exosome signal in the surrounding the presynaptic area is visible (**Fig 1A”’**, yellow line marks surrounding area in a radius of 1µm, corresponding to the postsynaptic area).

**Figure 1.**
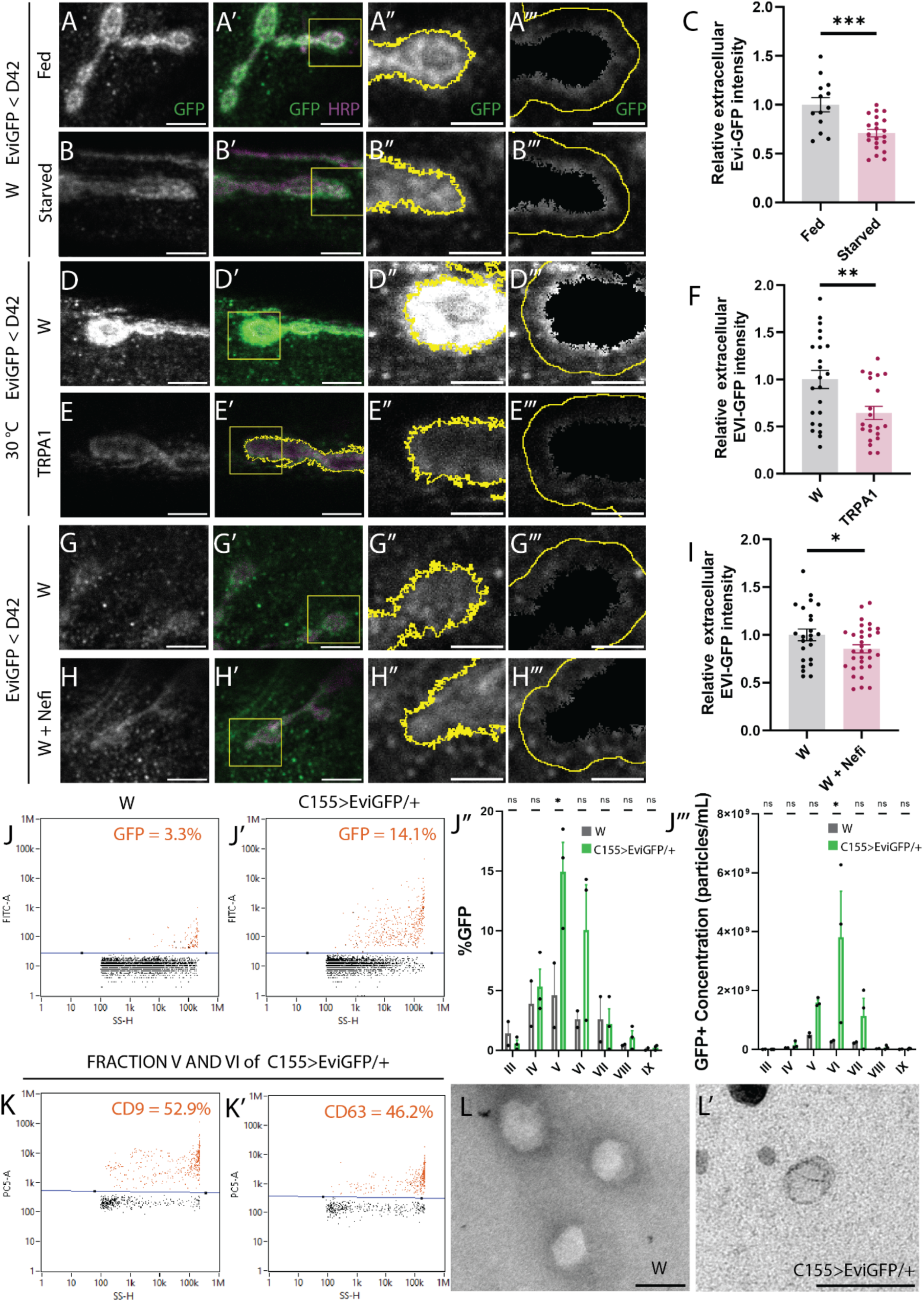
Autophagy induction decreases exosome release. (A, B, D, E, G, H) Representative images of neuromuscular junction (NMJ) boutons expressing EviGFP in grey. (A’, B’, D’, E’, G’, H’) The same representation but with EviGFP (green) and HRP (magenta) with the square in yellow that is going to be crop in the two subsequent columns. (A’’, B’’, D’’, E’’, G’’, H’’) The previous yellow square zoom in with EviGFP in grey and the silhouette of the HRP channel highlighted in yellow. (A’’’, B’’’, D’’’, E’’’, G’’’, H’’’) The same square with EviGFP in grey without the inside of the HRP silhouette. In yellow, it is that silhouette enlarged by 1 µm that is what is measured and represented in C, F, I when it is divided by the same area in HRP channel to control for laser power, antibody staining and macro. (A, A’, A’’, A’’’) Images of wildtype fed versus (B, B’, B’’, B’’’) starved NMJ and (C) its quantification that has been transformed dividing by the mean of wildtype fed. N (no. of animals) = 3, n (no. of images) = 13 (fed) and N = 5, n = 21 (starved) image fields per condition (p=0.0004). (D, D’, D’’, D’’’) Images of wildtype NMJ versus (E, E’, E’’, E’’’) the one expressing TRPA1 channel and (F) its quantification that has been transformed dividing by the mean of wildtype. N=5, n= 23 (W) and N = 5, n = 21 (TRPA1) image fields per genotype (p =0.0052). (G, G’, G’’, G’’’) Images of wildtype not treated and (H, H’, H’’, H’’’) treated with 1 μM Nefiracetam (Nefi) and (I) its quantification that has been transformed dividing by the mean of wildtype fed. N = 6, n = 24 (W) and N = 6, n = 32 (W + Nefi) image fields per condition (p = 0.0476). EviGFP is expressed using a motor neuron driver (D42-Gal4) Data was analyzed using student t-test (two-tailed) for normal distributions. Normality was analyzed with D’Agostino-Pearson Omnibus test * p<0.05, ** p<0,01, *** p<0.001, error bars = mean ± SEM. Scale bars represents 5 µm on left two columns and 2.5 µm on the right two columns. (J, J’) Dot plot of the events detected, FITC-A signal in Y as a function of the side scatter height (SS-H) in X, after setting a threshold for the noise, in orange is the GFP signal (J) in wildtype 3.3% and (J’) 14.1 % in C155>EviGFP/+. (J’’) Bar graphs representing the % of GFP+ events in fraction III-IX for the two different conditions: wildtype and C155>EviGFP/+. (J’’’) Bar graphs representing GFP+ events (particles) concentration in fractions III-IX expressed in particles/ml of two different conditions wild type and C155>EviGFP/+. Each dot in (J’’, J’’’) represent an independent experiment, n = 2 wild type (500 flies) and n = 3 in C155>EviGFP/+ (750 flies). The test for the analysis was Sidaks multiple comparison test and * p<0.05, error bars = mean ± SEM. (K, K’) Representative dot plots of the percentage of CD9+ (52.9%) (K) and CD63+ (46.2%) (K’) events displayed in orange from combined fractions V and VI from C155>EviGFP/+, PC5-A (APC) signal in y as a function of the side scatter height SS-H in X, after setting the threshold for the noise. (L, L’) Transmission electron microscopy (TEM) images of combined fractions V and VI in wild type (L) and C155>EviGFP/+ (L’). Scale bar represents 200 nm.

We first asked if induction of autophagy alters exosome release. Previously studies showed that autophagosomal formation at presynaptic terminals can be induced by different signals such as amino acid starvation and neuronal activity (Bademosi et al., 2023; Hernandez-Diaz et al., 2022; Soukup et al., 2016; Vanhauwaert, Kuenen, Masius, Bademosi, Manetsberger, Schoovaerts, Bounti, Gontcharenko, Swerts, Vilain, Picillo, Barone, Munshi, Vrij, Kushner, Gounko, Mandemakers, Bonifati, Meunier, Soukup, Verstreken, et al., 2017). We found that autophagy induction after amino-acid deprivation for a 4-hour period reduces exosome release compared to fed control animals (**Fig 1A-C).** We next tested if the induction of autophagy via neuronal activity affects exosome release similarly to amino acid starvation. Therefore, we employed the genetically encoded temperature-sensitive Drosophila TRPA1 (dTrpA1) cation channel to induce neuronal stimulation and activity. Specific expression and activation of dTrpA1 in motor neurons rapidly induces autophagy when flies are shifted to 30°C (Soukup et al., 2016). We used this assay in flies co-expressing the exosome marker EVI::GFP and dTrpA1 in motor neurons. Similar to our previous result, activation of autophagy by neuronal activation also reduces exosome release from the presynaptic compartment (**Fig 1D-F).** We further confirmed these results by using the drug Nefiracetam in combination with calcium that activates synaptic autophagy by opening L-type calcium channels (Bademosi et al., 2023; Nishizaki et al., 1998; Yoshii et al., 2000; Yoshii & Watabe, 1994). In line with our results, we observe that autophagy induction via Nefiracetam also leads to decrease exosome release at the NMJ synapse compared to untreated control animals (**Fig 1G-I).** Taken together our results indicate that autophagy and exosome release are connected at the synapse and that activation of autophagy via different stimulus (e.g. metabolic and neuronal activity) results in reduced levels of exosome release from the NMJ synapse.

To further validate our findings, we took advantage of an innovative fluorescent flow cytometry for nanoparticles (NanoFCM) approach to analyze fluorescent labelled exosomes from *Drosophila* adult brain neurons. With our settings the NanoFCM flow cytometer allows the visualization of particles between 30 to 200nm range and the simultaneous detection of fluorescent signals. To analyze specifically neuronal exosomes and to discriminate them from glial exosomes, we expressed EVI::GFP using a well-established Drosophila pan-neuronal driver (C155). We isolated exosomes from adult fly brains through a combination of ultracentrifugation and sucrose gradient. Analysis of extracted exosomes from wild type flies and flies expressing the EVI-GFP exosome marker in neurons showed that NanoFCM can specifically detect EVI-GFP positive neuronal exosomes (14,1% of the total signal) while in wildtype only GFP background signal was detected (**Fig 1 J, J’)**. We analyzed in more detail the different exosome-containing fractions and identified that fractions five and six have the highest amount of GFP positive exosomes (**Fig 1 J”, J”’).** Labelling of the exosome fractions five and six with antibodies against standard exosomal markers, such as CD63 and CD9 followed by analysis with the NanoFCM showed that the extracted exosomes are positive for both markers (**Fig 1 K, K’)**, confirming the identity of these extracellular vesicles as exosomes. Using transmission electron microscopy of this fraction we observe the typical shape of exosomes with a size below than 200 nm (**Fig 1 L, L’**), which ranks within the reported size of exosomes.

To understand how autophagy is connected to exosome release, we performed a candidate screen approach where we knock-down various autophagy core proteins functioning at different step in the autophagy pathway, following a step-wise targeted method from early phagosome formation, and quantified the variation of exosome release at the level of the synapse. We observe that knock down of Atg17/FIP200, a component of the ULK complex that acts at the initial nucleation step, leads to significant less exosome release (**Fig 2 A-B**). Likewise, knock down of Atg18, required for the expansion of the growing phagophore (Kotani et al., 2018), causes a significant reduction of exosome release (**Fig 2 C-D**). Closure of the growing phagophore depends on the autophagic protein Atg3 that mediated Atg8 conjugation to the autophagic membrane. We observe that also knock down of Atg3 leads to a significant reduction of exosome release (**Fig 2 E-F**). The autophagic protein Atg16 functions in a complex in the elongation of the phagophore and we observe that knock down of Atg16 leads to a significant reduction of exosome release (**Fig 2 G-H**). Once the autophagosome is formed, the autophagosome fuses with the lysosome to degrade the autophagic substrates. We were wondering if blocking the fusion of the autophagosome with the lysosome also effects exosome release. Chloroquine treatment has been shown to inhibits autophagic flux at the NMJ synapse by blocking the fusion of the autophagosome with the lysosomes at the Drosophila NMJ (Hernandez-Diaz et al., 2022). Interestingly, Chloroquine treatment did not affect exosome release at the NMJ synapse (**Fig 2 I-J**), indicating that autophagosome fusion with the lysosome does not affect exosome release. In contrast, our results may indicate that proteins functioning in the biogenesis of autophagosome could participate also during biogenesis of exosomes. Maturation of autophagosomes requires the fusion with late endosomes (Jäger et al., 2004; Takáts et al., 2013) and this is also true at the synapse, where autophagosomes fuse with late endosomes. Interestingly, multivesicular bodies (MVB), a specialized type of late endosomes, give rise to exosomes. Thus, both autophagy and exosome release mechanisms rely on the availability of late endosomes/MVBs. We reasoned that proteins associated to the MVB and to autophagosome biogenesis could be the molecular link between autophagy and exosome release and that molecules acting at this level could control the balance between autophagosome maturation and exosome formation. We used Pearson correlation to analyze the colocalization of different autophagic core proteins with Evi^GFP^ or Evi^mCherry^ positive puncta (corresponding to the MVBs that give rise to the exosomes). We detected no significant overlap of Atg8^mCherry^, Atg3^HA^, Atg16^HA^ and Atg17^GFP^ with the Evi marker (**Fig 2 K, L, M, N-and P**). However, we found significant co-localization of Atg18^GFP^ and Evi^mCherry^ positive puncta (MVBs) (**Fig 2 O and P**), suggesting that Atg18a may have a novel function during exosome release at the level of the MVBs.

**Figure 2.**
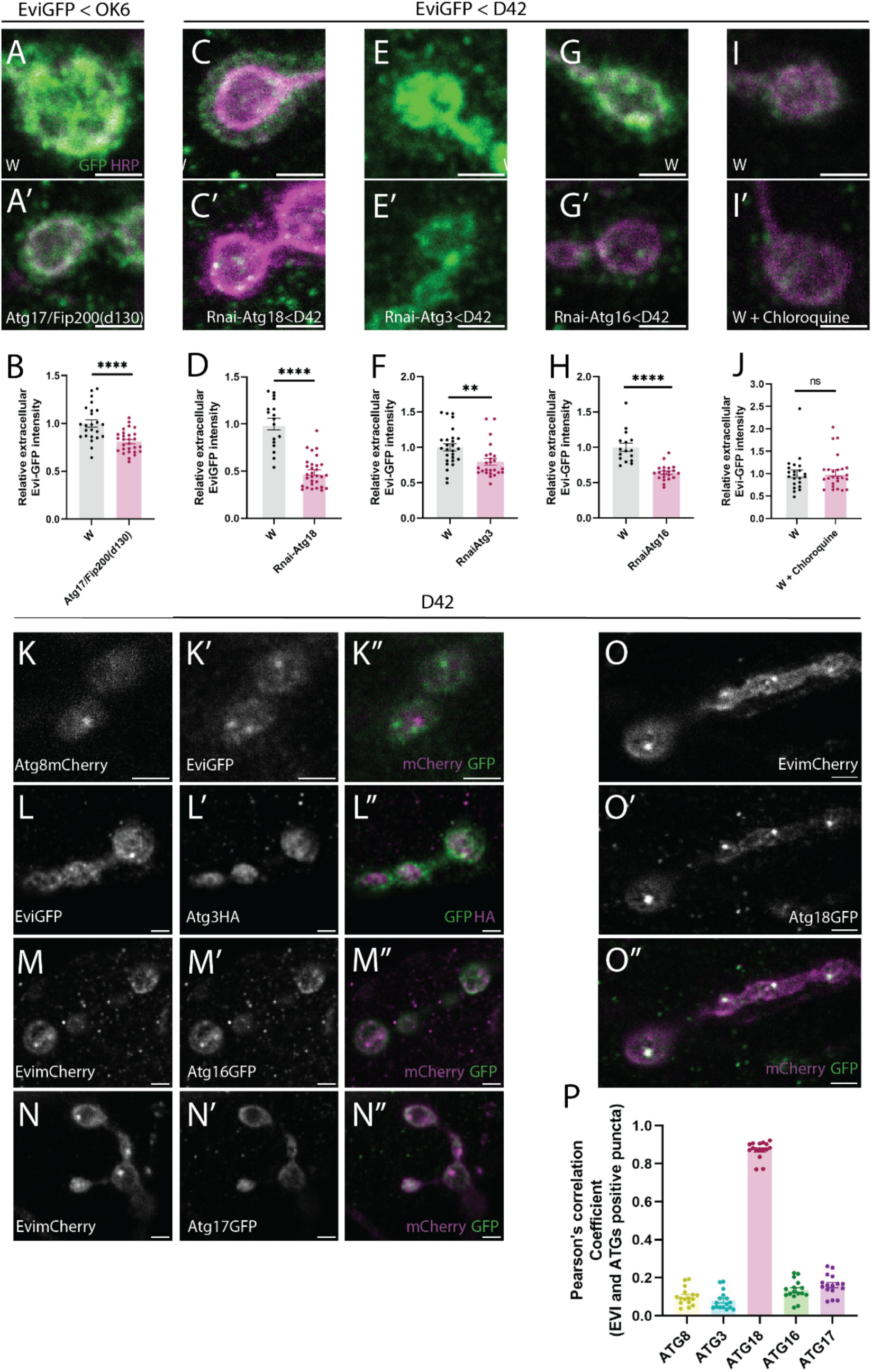
Atg18a is the protein involved in the reduction of the exosomes release. (A, A’, C, C’, E, E’, G, G’, I, I’) Representative images of neuromuscular junction (NMJ) boutons expressing EviGFP (green) and HRP (magenta). Scale bar represents 2.5 µm. (A) Image of wildtype versus (A’) the one expressing Atg17/Fip200d130 null mutant and (B) its quantification N = 5, n = 24 (W) and N = 5, n= 28 (Atg17/Fip200d130) p < 0.0001. EviGFP is expressed using a motor neuron driver (OK6-Gal4). (C) Image of wildtype versus (C’) the one expressing Rnai-Atg18a and (D) its quantification N = 5, n = 17 (W) and N = 9, n = 30 Rnai-Atg18a p < 0.0001. (E) Image of wildtype versus (E’) the one expressing Rnai-Atg3 and (F) its quantification. N = 6, n = 27 (W) and N = 5, n=25 Rnai-Atg3 p = 0.0017. (G) Image of wildtype versus (G’) the one expressing Rnai-Atg16 and (H) its quantification N = 3, n = 16 (W) and N = 8, n = 21 (Rnai-Atg16) p < 0.0001. (All the Rnai lines (C’) Atg18a, (E’) Atg3, (G’) Atg16, (are expressed under the motor neuron driver D42. (V) Image of wildtype not treated versus (I’) the one treated with chloroquine and (J) its quantification N = 5 n = 22 (W) and N = 5 n= 25 (W + Chloroquine) p = 0.9411. All the quantifications have been transformed dividing all the data by the mean of the wildtype of its respective control. EviGFP is expressed using a motor neuron driver (D42-Gal4). Data was analyzed using student t-test (two-tailed) for normal distributions and Mann Whitney test for not normal distribution. Normality was analyzed with D’agostino-Pearson Omnibus test ns p>0.05, * p<0.05, ** p<0,01, *** p<0.001, **** p<0.0001 error bars = mean ± SEM. (K, K’, K’’) Images of NMJ boutons expressing (K) atg8mCherry in grey, (K’) EviGFP in grey and (K’’) the merged in which Atg8mCherry is magenta and EviGFP is green. (L, L’, L’’) Images of NMJ boutons expressing (L) EviGFP in grey, (L’) Atg3HA in grey and (L’’) the merged in which EviGFP is green and Atg3HA is magenta. (M, M’, M’’) Images of NMJ boutons expressing (M) EvimCherry in grey, (M’) Atg16GFP in grey and (M’’) the merged in which EvimCherry is magenta and Atg16GFP is green. (N, N’, N’’) Images of NMJ boutons expressing (O) EvimCherry in grey, (N’) Atg17GFP in grey and (N’’) the merged in which EvimCherry is magenta and Atg17GFP is green. (O, O’, O’’) Images of NMJ boutons expressing (O) EvimCherry in grey, (O’) Atg18aGFP in grey and (O’’) the merged in which EvimCherry is magenta and Atg18aGFP is green. Scale bar represents 2.5 µm (P) Quantification of Pearson coefficient of Atg proteins (8, 3, 18, 16, 17 and 18) colocalization on Evi^Cherry^ or Evi^GFP^ puncta in the mask of Evi puncta (N = 6, n =16 Atg8; N = 5, n =16 Atg3; N = 5 n=16 Atg18a; N = 5 n= 16 Atg16; N = 5 n = 16 Atg17).

### Atg18a proteins levels are critical for exosome release

The autophagy core protein Atg18a (homologue to the human WIPI2) has been shown to be critical for autophagosomal formation (Proikas-Cezanne et al., 2015) and at the presynaptic terminal Atg18a function is required for maturation of the autophagosome (Vanhauwaert, Kuenen, Masius, Bademosi, Manetsberger, Schoovaerts, Bounti, Gontcharenko, Swerts, Vilain, Picillo, Barone, Munshi, Vrij, Kushner, Gounko, Mandemakers, Bonifati, Meunier, Soukup, Verstreken, et al., 2017). To further confirm that Atg18a has an additional function in exosome release at the synapse we analyzed Atg18 null mutant flies and, similar to Atg18a knock down, loss of Atg18a (w;; atg18a^-/-^) leads to significant reduction of exosome release from the neuronal synapse (**Fig 3 A-C**). Consequently, overexpression of Atg18a (w;;Atg18a^HA^) in motor neurons leads to increased exosome release from the synapse (**Fig 3 D-F**). Our data point out that Atg18a is a positive regulator of exosome release and that Atg18a proteins levels are critical for the biogenesis of exosomes.

**Figure 3.**
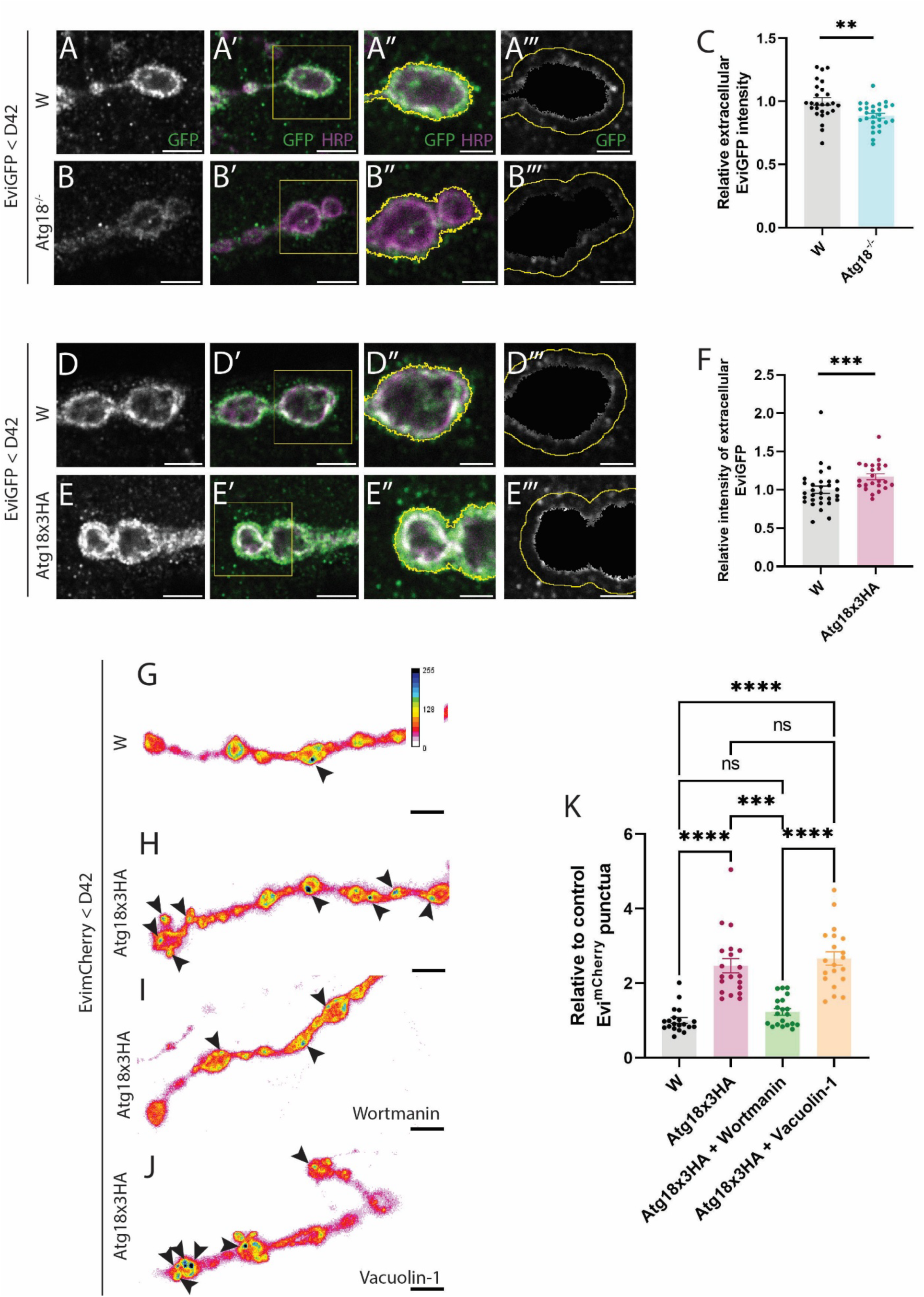
Atg18a modulates exosomes release and does it through PI3P and PI(3,5)P_2_. (A, B, D, E) Representative images of neuromuscular junction (NMJ) boutons expressing EviGFP in grey. (A’, B’, D’, E’) The same representation but with EviGFP (green) and HRP (magenta) with the square in yellow that is going to be crop in the two subsequent columns. (A’’, B’’, D’’, E’’) The previous yellow square zoom in with EviGFP in grey and the silhouette of the HRP channel highlighted in yellow. (A’’’, B’’’, D’’’, E’’’) The same square with EviGFP in grey without the inside of the HRP silhouette. In yellow, it is that silhouette enlarged by 1 µm that is what is measured and represented in C, F when it is divided by the same area in HRP channel to control for laser power, antibody staining and macro. (A, A’, A’’, A’’’) Images of wildtype versus (B, B’, B’’, B’’’) Atg18a^-/-^ NMJ and (C) its quantification that has been transformed dividing by the mean of wildtype. N = 5, n =26 (W) and N = 5, n = 27 (Atg18a^-/-^) image fields per condition (p = 0.0020). (D, D’, D’’, D’’’) Images of wildtype NMJ versus (E, E’, E’’, E’’’) Atg18ax3HA (overexpression) and (F) its quantification that has been transformed dividing by the mean of wildtype. N=6, n= 30 (W) and N = 4, n =23 (Atg18ax3HA) image fields per genotype (p =0.0009). EviGFP is expressed using a motor neuron driver (D42-Gal4) Data was analyzed using student t-test (two-tailed) for normal distributions. Normality was analyzed with D’Agostino-Pearson Omnibus test ** p<0,01, *** p<0.001, error bars = mean ± SEM. Scale bars represents 5 µm on left two columns and 2.5 µm on the right two columns. (G, H, I, J) Images of NMJ boutons of wild type (G), Atg18ax3HA (H), Atg18ax3HA with 5uM Wortmanin, a PI3K inhibitor (I) and Atg18ax3HA with 5uM Vacuolin-1 a PIKfyve inhibitor (J). Fluorescence intensities shown using indicated scale in G. Arrows indicate the number of EvimCherry dots (multivesicular bodies). Scale bars are 5 µm (K) Quantification of the EvimCherry dots coming from G, H, I, J. n = 20 N=5 in all conditions that has been divided by the mean of wild type condition. Normality was analyzed with D’Agostino-Pearson Omnibus test. ANOVA Kruskal-Wallis test and a Dunn’s test was used. Significance of statistical difference was defined as **** =p<0.0001, *** =p<0.001, ns= p>0.05.

### Atg18a functions in the biogenesis of multivesicular bodies

The biogenesis of MVBs is essential for exosome release and the number of MVBs that fuse with the plasma membrane determines the quantity of exosome in the extracellular space (Lauwers et al., 2018). Thus, we were asking if Atg18a plays a role in MVB biogenesis or facilitates the fusion of MVBs with the plasma membrane.

Using confocal microscopy, we observed that overexpression of Atg18a leads to increased number of Evi^mCherry^ positive MVBs (**Fig 3 G,H and K**). Drosophila Atg18a and the mammalian homologue WIPI2 are folded into a seven-bladed β-propeller that allow Atg18 to bind to membrane lipids of preautophagosomal membranes or other organelles by specifically binding to the phosphoinositides PI3P and PI(3,5)P2 (Proikas-Cezanne et al., 2015). Intriguingly, Atg18 binding to PI3P promotes autophagy and evidence indicate that proteins belonging to the WIPI protein family can also bind endosomal membranes via the interaction to PI(3,5)P2 (De Leo et al., 2021). To further understand the molecular function of Atg18a in exosome release we wanted to uncover if Atg18a function in exosome release depends on the recognition of PI3P and PI(3,5)P2 on MVB membranes. For this purpose, we reduce the levels of those phosphoinositides by inhibiting the PI3P producing enzyme Vps34 and the PI(3,5)P2 producing enzyme Pikfyve by feeding the animals with the drug Wortmanin or Vacuolin-1 respectively (Scott et al., 2004)(Vanhauwaert, Kuenen, Masius, Bademosi, Manetsberger, Schoovaerts, Bounti, Gontcharenko, Swerts, Vilain, Picillo, Barone, Munshi, Vrij, Kushner, Gounko, Mandemakers, Bonifati, Meunier, Soukup, Verstreken, et al., 2017). We observed that the treatment with Wortmanin in flies overexpressing Atg18a leads to a reduction of Evi^mCherry^ positive puncta comparable to wildtype animals (**Fig 3 I, K**). In contrast treatment with Vacuolin-1 does not significantly decrease Evi^mCherry^ positive puncta in those flies (**Fig 3 J, K**). Taken together, these results indicate that Atg18a function *via* PI3P binding is required for biogenesis of Evi positive MVBs.

### Exosome release is altered by a Parkinson disease mutation in Synaptojanin

Previous studies showed that Atg18a/WIPI2 is connected to the presynaptic protein Synaptojanin (Synj) (Vanhauwaert, Kuenen, Masius, Bademosi, Manetsberger, Schoovaerts, Bounti, Gontcharenko, Swerts, Vilain, Picillo, Barone, Munshi, Vrij, Kushner, Gounko, Mandemakers, Bonifati, Meunier, Soukup, Verstreken, et al., 2017). Mutations in the human homologue SYNJ1 cause Parkinson’s disease (Krebs et al., 2013). At the NMJ synapse, Synj/SYNJ1 function *via* Atg18a/WIPI2 is critical for autophagosomal formation and the expression of Synj harboring the Parkinson disease causative mutation R258Q results in a blockage in autophagosome maturation that generates an accumulation of Atg18a positive immature autophagosomes (Vanhauwaert, Kuenen, Masius, Bademosi, Manetsberger, Schoovaerts, Bounti, Gontcharenko, Swerts, Vilain, Picillo, Barone, Munshi, Vrij, Kushner, Gounko, Mandemakers, Bonifati, Meunier, Soukup, Verstreken, et al., 2017). Thus, we asked whether the Parkinson disease mutation of Synj also affects exosome release. We analyzed the effect of pathological Synj^R258Q^ (Synj^RQ^) on exosome release in neurons of adult brains using the NanoFCM approach by quantifying EVI^GFP^ fluorescent labelled exosomes from Drosophila adult brain neurons. We observed that isolated exosomes from Synj^RQ^ (C155>Evi-GFP, Synj^RQ^), and control (C155>Evi-GFP) brains have comparable morphology and size distribution in electron micrographs (**Fig 4 A, A’)** and NanoFCM flow cytometer (**Fig 4 C, C’).** Interestingly, NanoFCM measurements show a significant decrease of Evi^GFP^ positive exosomes in Synj^RQ^ flies compared to control flies (**Fig 4 B, D).** This reduction was not caused by a size variation of Evi^GFP^ positive exosome from Synj^RQ^ control brains (**Fig 4 E).** Analysis of the different exosome fraction showed that Evi^GFP^ positive exosome isolated from Synj^RQ^ control brains separate similarly and concentrate around fraction six (**Fig 4 F)** and there were no significant differences in exosomes size in the different fractions (**Fig 4 G).** Taken together our result show that exosome release in Synj^RQ^ mutant brain is decreased and this is not caused by any changes in morphology or size of exosomes.

**Figure 4.**
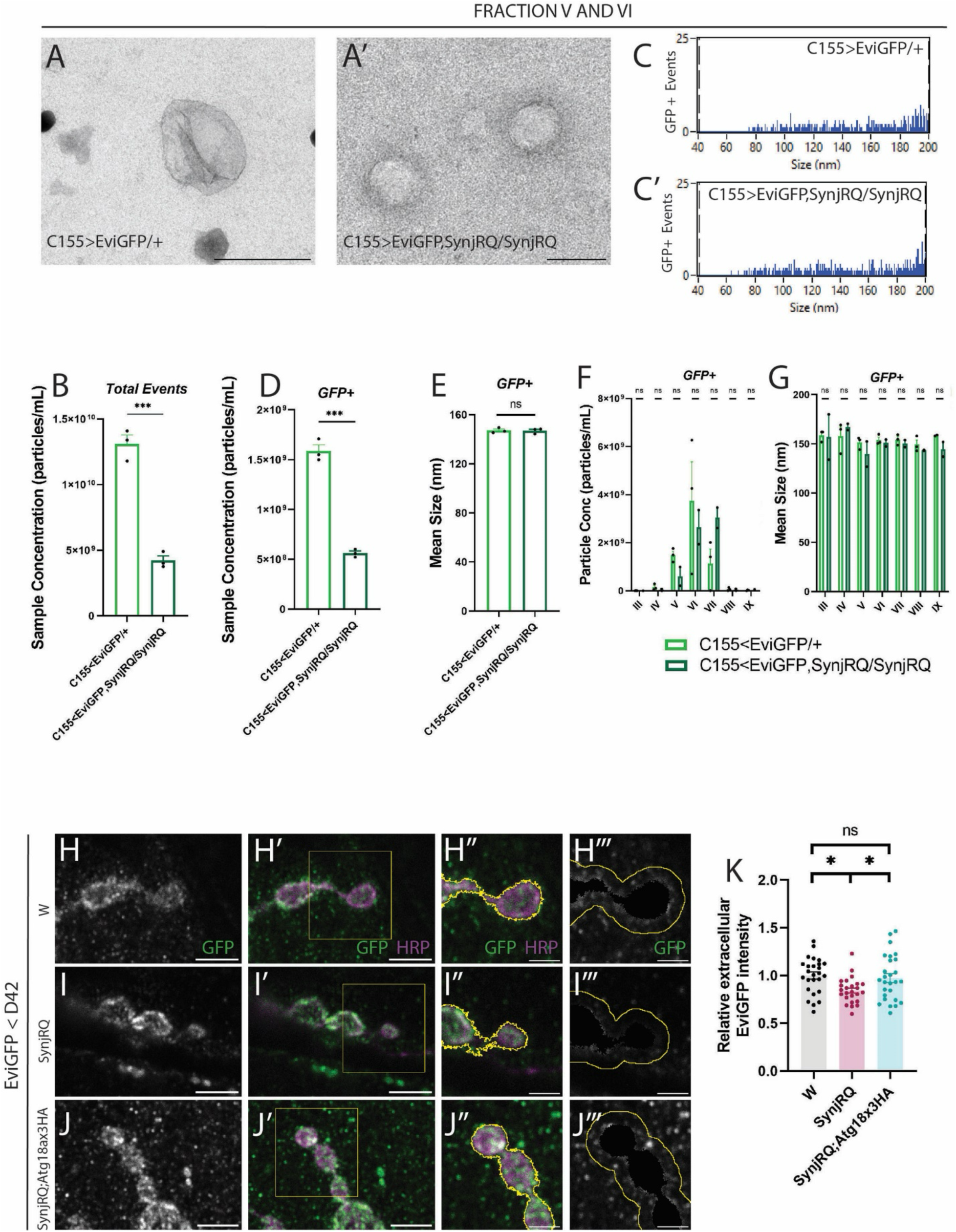
Synj^RQ^ mutation decreases exosome release and overexpression of Atg18a can rescue the phenotype independent of its autophagy function. (A, A’) Transmission electron microscopy (TEM) images of combined fractions V and VI in (A) C155>EviGFP/+ and (A’) C155>EviGFP,SynjRQ/SynjRQ. Scale bar represents 200 nm. (B) Bar graphs representing the sample concentration in particles/ml of two different conditions C155>EviGFP/+ and C155>EviGFP,SynjRQ/SynjRQ. (C, C’) Bar graphs representing the size distribution of GFP+ positive events (exosomes) in (C) C155>EviGFP/+ and (C’) C155>EviGFP,SynjRQ/SynjRQ. (D) Bar graphs representing the sample concentration expressed in particles/ml of GFP+ particles (exosomes) in C155>EviGFP/+ and C155>EviGFP,SynjRQ/SynjRQ. (E) Bar graphs representing the mean size of GFP+ particles (corresponding to exosomes) in C155>EviGFP/+ and C155>EviGFP,SynjRQ/SynjRQ. (F) Bar graphs representing the particle concentration expressed in particles/ml of GFP+ particles (exosomes) in fractions III-IX of C155>EviGFP/+ and C155>EviGFP,SynjRQ/SynjRQ. (G) Bar graphs representing the mean size in nm of GFP+ particles (exosomes) in the different fractions of C155>EviGFP/+ and C155>EviGFP,SynjRQ/SynjRQ. Mixed effects analysis with Sidak’s multiple comparison test and unpaired t-test were used. (H, I, J) Representative images of neuromuscular junction (NMJ) boutons expressing EviGFP in grey. (H’, I’, J’) The same representation but with EviGFP (green) and HRP (magenta) with the square in yellow that is going to be crop in the two subsequent columns. (H’’, I’’, J’’) The previous yellow square zoom in with EviGFP in grey and the silhouette of the HRP channel highlighted in yellow. (H’’’, I’’’, J’’’) The same square with EviGFP in grey without the inside of the HRP silhouette. In yellow, it is that silhouette enlarged by 1 µm that is what is measured and represented in K when it is divided by the same area in HRP channel to control for laser power, antibody staining and macro. (H, H’, H’’, H’’’) Images of wildtype versus (I, I’, I’’, I’’’) SynjRQ, (J, J’, J’’, J’’’) SynjRQ;Atg18ax3HA and (K) its quantification that has been transformed dividing by the mean of wildtype. EviGFP is expressed using a motor neuron driver (D42-Gal4) N = 5, n = 26 (W) and N = 5, n = 25 (SynjRQ) N = 5, n = 27 (SynjRQ;Atg18ax3HA) image fields per condition. Normality was analyzed with D’Agostino-Pearson Omnibus test. ANOVA Ordinary test for nomal distribution was used and a Tukey test was used. Data are expressed as mean ± SEM. Significance of statistical difference was defined as ***p<0.001, * =p<0.05, ns= p>0.05.

To confirm if Synj^RQ^ alters exosome release from presynaptic terminals, we analyzed exosome release by confocal microscopy at the NMJ synapse. We observe that similarly to our NanoFCM analysis, Synj^RQ^ mutant animals release significant less exosomes from the presynaptic terminal (**Fig 4, H-I, K).** Since overexpression of Atg18a increases exosome release, we tested if overexpression of Atg18a could rescue altered exosome release observed in Synj^RQ^ mutant animals. Using confocal microscopy, we revealed that overexpression of Atg18a in Synj^RQ^ mutant animals (Synj^RQ^; Atg18a^HA^) restores exosome release to levels comparable to wildtype flies (**Fig 4, J-K)**.

### Overexpression of Atg18a rescues Synj^RQ^ associated locomotor dysfunction

Parkinson disease is typically diagnosed when motor symptoms appear, and at that time around 30%-70% of the dopaminergic (DA) neurons in the substantia nigra pars compacta (SNc) have degenerated (Fearnley and Lees, 1991; Cheng et al., 2010; Giguère et al., 2018). To further understand how alterations in exosome release contribute to neuronal dysfunction and the onset of neurodegeneration, we decided to investigate how Atg18a overexpression may affect locomotion, a major dysfunction associated to Parkinson disease. To analyse locomotion, we performed a negative geotaxis assay that has been previously validated in *Drosophila* models of disease that present locomotion deficits (Feany & Bender, 2000). As expected, Synj^RQ^ mutant animals have significant reduced locomotion compared to control animals (**Fig 5, A)**. Interestingly, locomotion defects in Synj^RQ^ can be rescued by the solely overexpression of Atg18a in dopaminergic neurons (Synj^RQ^; Atg18a^HA^< TH-Gal4). Patients with Parkinson mutation R258Q in SYNJ1 show, in addition to motor symptoms, epileptic seizures (Hardies et al., 2016; Krebs et al., 2013; Quadri et al., 2013). Studies in Drosophila report the existence of stress-sensitive seizure-like behaviors and paralysis mutations upon mechanical stimulation or “bang assay” (Fergestad et al., 2006). We thus used the bang sensitivity assay to analyze if mechanical induced seizure-like behaviors were also observed in Synj^RQ^ mutant animals. Interestingly, animals expressing the Synj^RQ^ PD mutation have significant longer recovery time after the induction of seizure-like behaviors s compared to control animals (**Fig 5, B)**. Furthermore, overexpression of Atg18a in dopaminergic neurons was able to rescue the seizure-like behavior observed in Synj^RQ^ mutant animals, as those flies have a recovery that is statistically not different from control flies (**Fig 5B)**. We further performed a hybrid seizure-locomotion assay combining the bang and the negative geotaxis assay to quantify sensitivity to seizure and locomotion performance (Tao et al., 2011). Our results show that induction of seizures greatly delay the initiation of locomotion in Synj^RQ^ flies compared to control flies and overexpression of Atg18a in dopamine neurons of Synj^RQ^ mutant flies also improves locomotion after seizure recovery (**Fig 5B’**). Taken together our results show that Synj^RQ^ mutant’s animals display locomotion defects and seizure-like behavior and that these defects can be restored by the overexpression of Atg18a in dopaminergic neurons, linking Atg18a alterations to core symptoms of Parkinson’s disease. A hallmark of Parkinson disease is the loss of dopaminergic neurons. Drosophila is an excellent model to study dopaminergic neuron loss since dopaminergic neurons in the adult brain are organized in anatomical clusters (**Fig 5C**). Moreover, we previously showed that dopaminergic neuron degeneration of the dopaminergic neuron clusters PPM3 and PPM1 in Synj^RQ^ mutant flies (Vanhauwert et al. 2017). Here we show that dopaminergic neuron loss of cluster PPM3 and PPL1 in 10d old Synj^RQ^ can be rescued by overexpression of Atg18 in dopaminergic neurons (**Fig 5C’,C”**). We then tested if overexpression of Atg18a was restoring Synj^RQ^ dysfunction through exosome release or autophagy. Previous studies showed that synaptic autophagy induction is blocked in Synj^RQ^ mutant animals (Vanhauwaert, Kuenen, Masius, Bademosi, Manetsberger, Schoovaerts, Bounti, Gontcharenko, Swerts, Vilain, Picillo, Barone, Munshi, Vrij, Kushner, Gounko, Mandemakers, Bonifati, Meunier, Soukup, Verstreken, et al., 2017). To monitor autophagy, we expressed the autophagy marker Atg8^mCherry^ in motor neurons (Soukup et al., 2016). Under basal conditions only few autophagosome (Atg8^mCherry^ positive dots, arrows) are observed, but overexpression of Atg18a leads to significant increased autophagy levels (**Fig 5, D, D”’, E)**. However, we did not observe that Atg18a overexpression increases autophagy in Synj^RQ^ mutant animals **(Fig 5, D, D’, D’’, D’’’, E)**. These results indicate that although Atg18a overexpression induce autophagy and increases exosome release in wild type flies, this is not the case in the context of the Synj1^RQ^ PD mutant flies, where the overexpression of Atg18a only restore exosome release. Hence, our results indicate that increasing exosome release by overexpressing Atg18a is sufficient to restore PD associated dysfunctions in Synj^RQ^ mutant animals.

**Figure 5.**
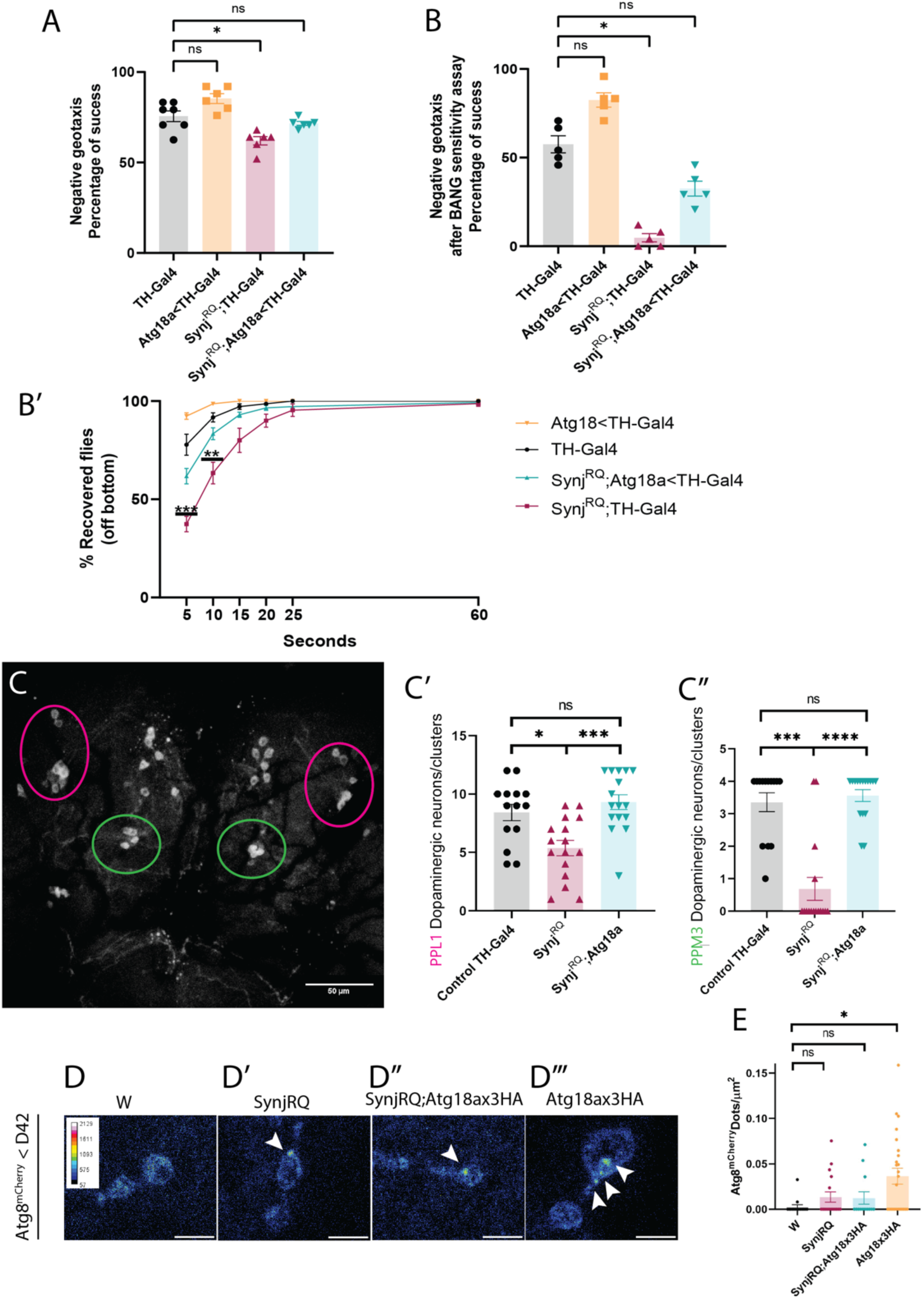
Synj^RQ^ mutation decreases movement behavior, predispose to seizures and neurodegeneration; and overexpression of Atg18a can rescue the phenotype independent of its autophagy function. (A) Quantification of the success in % of the negative geotaxis assay (also called the climbing assay) of 10 days old flies Each dot represents n = females and males (TH-Gal4 (n = 7), Atg18×3HA/TH-Gal4 (n = 6), SynjRQ;TH-Gal4 (n = 6), SynjRQ;Atg18×3HA/TH-Gal4(n = 6)). (B) Quantification of the success in % of the negative geotaxis assay (also called the climbing assay) of 10 days old flies after the bang sensitivity assay Each dot represents n = females and males (TH-Gal4 (n = 5), Atg18×3HA/TH-Gal4 (n = 5), SynjRQ;TH-Gal4 (n = 5), SynjRQ;Atg18×3HA/TH-Gal4 (n = 5)) Normality was analyzed with D’Agostino-Pearson Omnibus test. ANOVA Kruskal-Wallis test and a Dunn’s test was used. Significance of statistical difference was defined as * =p<0.05, ns= p>0.05. (B’) Graph representing the recovery off the bottom in comparison of the four genotypes 0.5 s after the bang assay Each dot represents n = females and males (TH-Gal4 (n = 5), Atg18×3HA/TH-Gal4 (n = 5), SynjRQ;TH-Gal4 (n = 5), SynjRQ;Atg18×3HA/TH-Gal4 (n = 5)) Normality was analyzed with D’Agostino-Pearson Omnibus test. 2way ANOVA was used pot-hoc test. Significance of statistical difference was defined as ** =p<0.01, *** =p<0.005, ns= p>0.05. (C) Representative image of anti-TH staining (to mark dopaminergic neuron clusters) in the posterior side of fly brain. Note the PPM3 and PPL1 dopmainergic neuron clusters are marked in green and magenta respectively. (C’) Quantification of the number of TH positive dopaminergic neurons in PPM3 clusters in 10-days old flies of the indicated genotypes. (C”) Quantification of the number of TH positive dopaminergic neurons in PPL1 clusters in 10-days old flies of the indicated genotypes. Statistical analysis of dopaminergic neurons shows fewer neurons in PPM3 and PPL1 in Synj^RQ^ mutant flies compared to control flies and Synj^RQ^ mutant flies overexpressing Atg18 in dopaminergic neurons (n>8 adult *Drosophila* brains per genotype).(D, D’, D’’, D’’’) Images of NMJ boutons of wild type (D), SynjRQ (D’), SynjRQ;Atg18ax3HA (D’’), Atg18ax3HA (D’’’) expressing Atg8mCherry. Fluorescence intensities shown using indicated scale in D. Arrows indicate the number of Atg8mCherry dots (autophagosomes). Scale bars are 2.5 µm (E) Quantification of the Atg8mCherry dots coming from (D, D’, D’’, D’’’). N = 3, n = 15 (W); N = 5, n = 17 (SynjRQ); N = 4, n = 13 (SynjRQ;Atg18×3HA); N =5, n = 25 (Atg18×3HA). that has been divided by the mean of wild type condition. Normality was analyzed with D’Agostino-Pearson Omnibus test. ANOVA Kruskal-Wallis test and a Dunn’s test was used. Significance of statistical difference was defined as * =p<0.05, ns= p>0.05.

## DISCUSSION

Exosomes participate in intercellular communication in the nervous system supporting, like autophagy, neuronal function, development and even disposal of pathogenic proteins. Interestingly, some stimuli regulating autophagy, such as nutrient deficiency, calcium concentration or even neuronal activity seem to also regulate exosome release. Presynaptic terminals are very dynamic compartments and rely on continues turnover over synaptic components to maintain neurotransmission. Therefore, correct functioning on efficient protein sorting and recycling is essential to maintain synaptic protein homeostasis and defects in synaptic homeostasis could result in synaptic dysfunction. Together with the observation that synaptic dysfunction precedes neuronal loss in many neurodegenerative diseases, dysfunction of the process regulation synaptic proteostasis could be at the root of neurodegeneration. Autophagy as well as exosome release at the presynaptic terminal has tightly linked to the sorting and degradation of synaptic components (Blanchette et al., 2022)(Kiral et al., 2020). Thus, clarifying the function and regulation of autophagy and exosome release at the presynaptic terminal is essential to understand how these processes participate in synaptic function, neurodegeneration and brain pathology. Unfortunately, opposite effects of autophagy induction in the process of exosome release are described in the scientific literature, suggesting that the crosstalk between these pathways is a complex and context dependent process. Here, we have deciphered how autophagy regulates exosome release *in vivo* at the presynaptic compartment. Our work shows that, at the synapse, activation of autophagy results in a decrease of exosome release. This was true for both autophagy induction *via* amino acid starvation and neuronal activity, indicating that the interplay between these two pathways could be mediated by core mechanisms functioning in autophagy and not *via* specific regulators. To further understand this interplay, we also tested how blocking autophagy at different steps affected exosome release. Evidence reports that inhibition of the acidification of the lysosome, a crucial step in autophagy that mediates the fusion of the autophagosome with the lysosome, can increase the secretion of APP *via* exosomes (Miranda et al., 2018). Thus, we wonder if blocking this final step in autophagy could results in higher number of released exosomes from the synapse. However, when we abolished lysosomal acidification and the formation of the autolysosome using the drug chloroquine, we detected no significant change in the number of exosomes secreted from the synaptic compartment. This indicates that, at least under physiological conditions at the presynaptic terminal, blocking the fusion of the autophagsosome with the lysosome does not affect exosome release. Next, we sought to investigate how disruptions in the initial steps of the autophagy pathway, the biogenesis of the autophagosome, affected exosome release. The autophagic core proteins Atg17, Atg18, Atg3 and Atg16 participate in different steps of autophagosome biogenesis and reduction of these proteins also results in a reduction of exosome release from the presynaptic compartment. This indicates that early steps of the autophagy pathway like initiation and nucleation of the autophagosome are crucial for exosome biogenesis or release. These results further suggest that alteration of autophagy at the presynaptic terminal could be connected or contribute to defective exosome release. The processes of autophagy and exosome release share the MVB as a common organelle, and MVBs can be found not only in the neuronal soma but also at the synapse. Interestingly, from the autophagic core proteins we tested to affect exosome release, only the autophagy protein ATG18a/WIPI2 localize with Evi^GFP^ positive MVBs, suggesting that this protein could function directly in both pathways. Accordingly, overexpression of Atg18a increase the number of exosomes released from the synapse and also leads to an increase in the number of synaptic MVBs, further suggesting that Atg18a regulates exosome release at this level. Atg18a is a PROPPIN (β-propeller proteins that bind phosphoinositides) family protein, that are known to have specific binding sites for the PI(3)P or PI(3,5)P2 phosphoinositides. Inhibition of PI3K, a kinase that regulates membrane trafficking and catalyzes the production of PI(3))P, reduces the excess of MVBs generated in response to Atg18a overexpression, while inhibition of PIKfyve, a kinase that produces PI(3,5)P2 has no effect in MVB formation. These results indicates that binding of Atg18a primarily to PI(3)P and not PI(3,5)P is necessary for Atg18a function in exosome biogenesis. PI(3)P levels have been shown to be critical for the recruitment of members of the ESCRT complex to the MVB and the formation of intraluminal vesicle (Wenzel et al., 2018). This is particular interesting since the generation of intraluminal vesicle is an essential step in exosome biogenesis. Our work gives also further insights into the autophagy exosome interplay in disease conditions. We report here that synapses harboring a PD mutation in the protein Synj show reduced exosome release. Previous studies demonstrated that autophagy is impaired in Synj PD mutant synapses, as this mutation restrains the phosphatase activity of Synj SAC1 domain (Vanhauwaert, Kuenen, Masius, Bademosi, Manetsberger, Schoovaerts, Bounti, Gontcharenko, Swerts, Vilain, Picillo, Barone, Munshi, Vrij, Kushner, Gounko, Mandemakers, Bonifati, Meunier, Soukup, Verstreken, et al., 2017). As a result of the defective phosphatase activity of Synj ^RQ^ on the phosphoinositols PI3P and PI(3,5)P_2_, the protein Atg18a cannot be released from the nascent autophagosomes and the process of autophagosome biogenesis is blocked. Our results show that Atg18a overexpression increases autophagy induction as well as exosome release in wild type synapses, suggesting that the availability of Atg18a regulates both processes in this compartment. Our results suggests that in Synj PD mutant synapses, Atg18a is a limiting factor in exosome biogenesis and those immature autophagosomes could act as an organelle trap preventing this protein to function in exosome release. Accordingly, we show that overexpression of Atg18a in Synj PD mutant synapses restores exosome release without rescuing autophagy induction. Patients with the Synj ^RQ^ mutation undergo early-onset Parkinsonism (EOP) with epileptic seizures (Hardies et al., 2016; Krebs et al., 2013; Quadri et al., 2013). The locomotion deficit and seizures associated to these patients has been further confirmed in mice models harboring the human Synaptojanin1 (SYNJ1^R258Q^) mutation (Cao et al., 2017) and we further validated that this mutation also induces locomotion defects and higher sensitivity and longer recovery from seizure-like behavior in our *Drosophila* model. Unexpectedly, rescuing the role of Atg18a in exosome release in dopaminergic neurons seems sufficient to rescue the locomotion deficits exhibited in Synj mutant PD *Drosophila* animals, indicating that exosomes release plays crucial roles in neuronal dopaminergic function and suggesting that defects in autophagy could also worsen disease progression by deregulating exosome release. Furthermore, these results indicate that exosome release from dopaminergic neurons is required to maintain proper locomotion, and impaired exosome release results in defective locomotion and seizure-like behavior that are also observed in patients with Synj PD mutations. Interestingly, dopaminergic neurodegeneration observed in Synj PD mutant flies can also be rescued by Atg18a overexpression indicating that reestablishing exosome release in dopaminergic neurons is sufficient to prevent neurodegeneration of these neurons potentially as compensatory mechanisms. Interestingly, under physiological conditions, exosome release from neurons has been shown to support the formation and activity of neural circuits (Lee et al., 2018)(Sharma et al., 2019). Currently, Parkinson associated symptoms are typically treated with L-DOPA. However, this treatment leads only to an alleviation of the symptoms and becomes more ineffective with the progression of the disease. Furthermore, not all PD patients respond positively to this treatment and for instance patients with the SYNJ1 ^RQ^ mutation respond poorly to L-DOPA. Therefore, alternative treatment that could alleviated symptoms and potentially support neuronal viability and or dopaminergic circuit activity would be a further step forward to effective treatment of the disease. However, further studies are required to establish the function of Atg18a/WIPI2 in dopaminergic circuits and to understand the precise molecular function of Atg18a/WIPI2 before developing novel drugs that targets WIPI2 function.

## MATERIALS AND METHODS

### Fly Stocks, genetics and housing

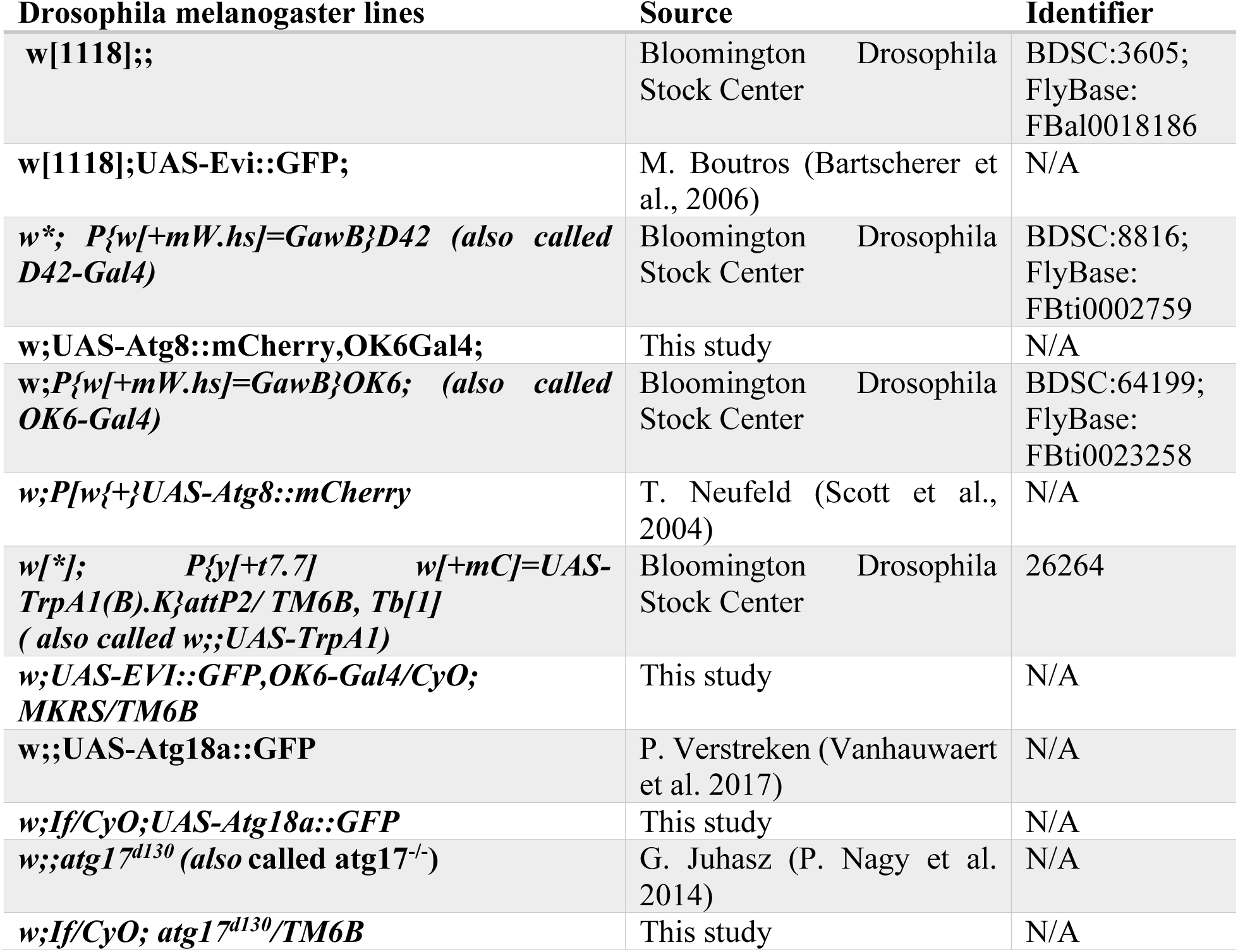
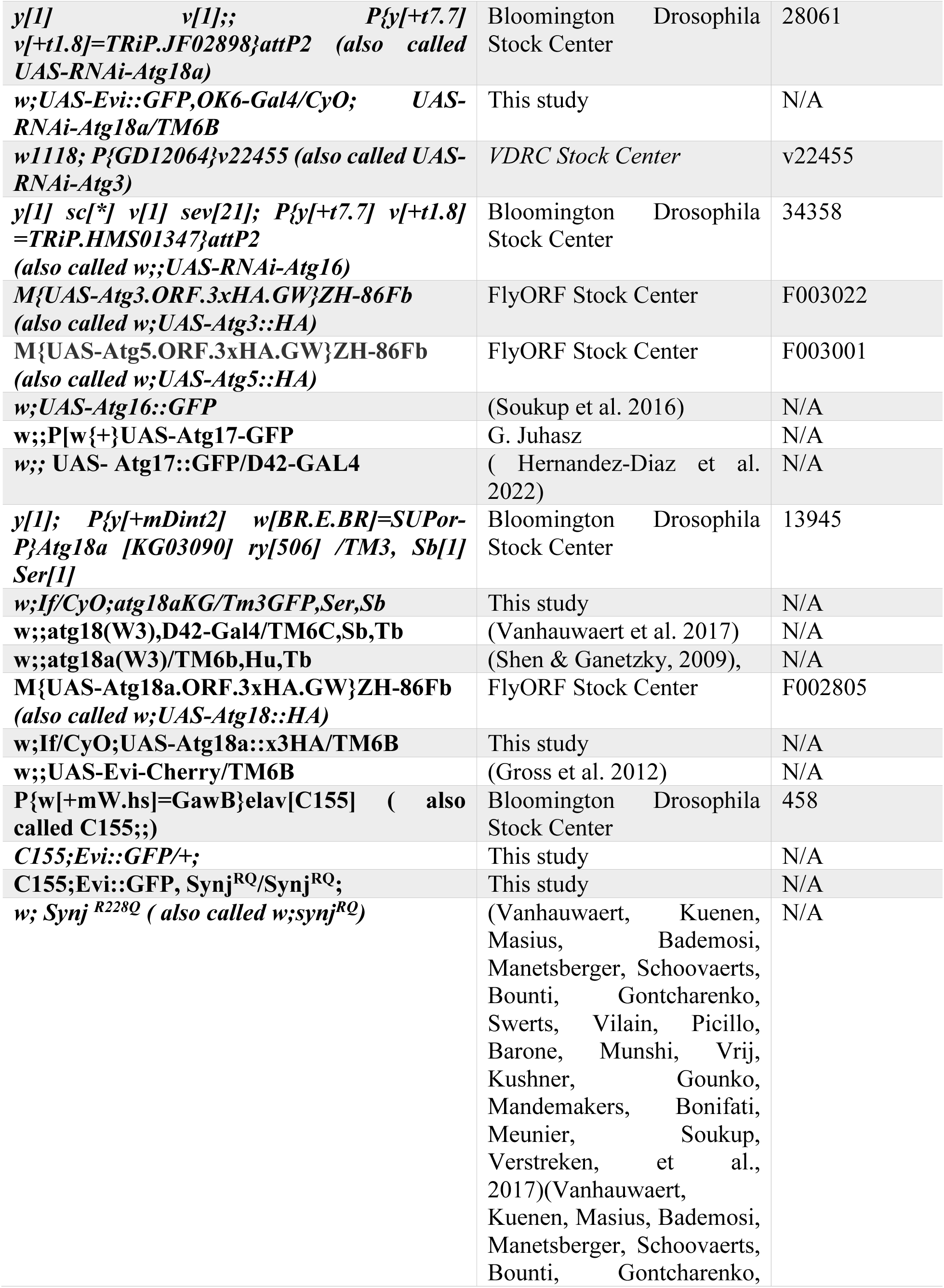
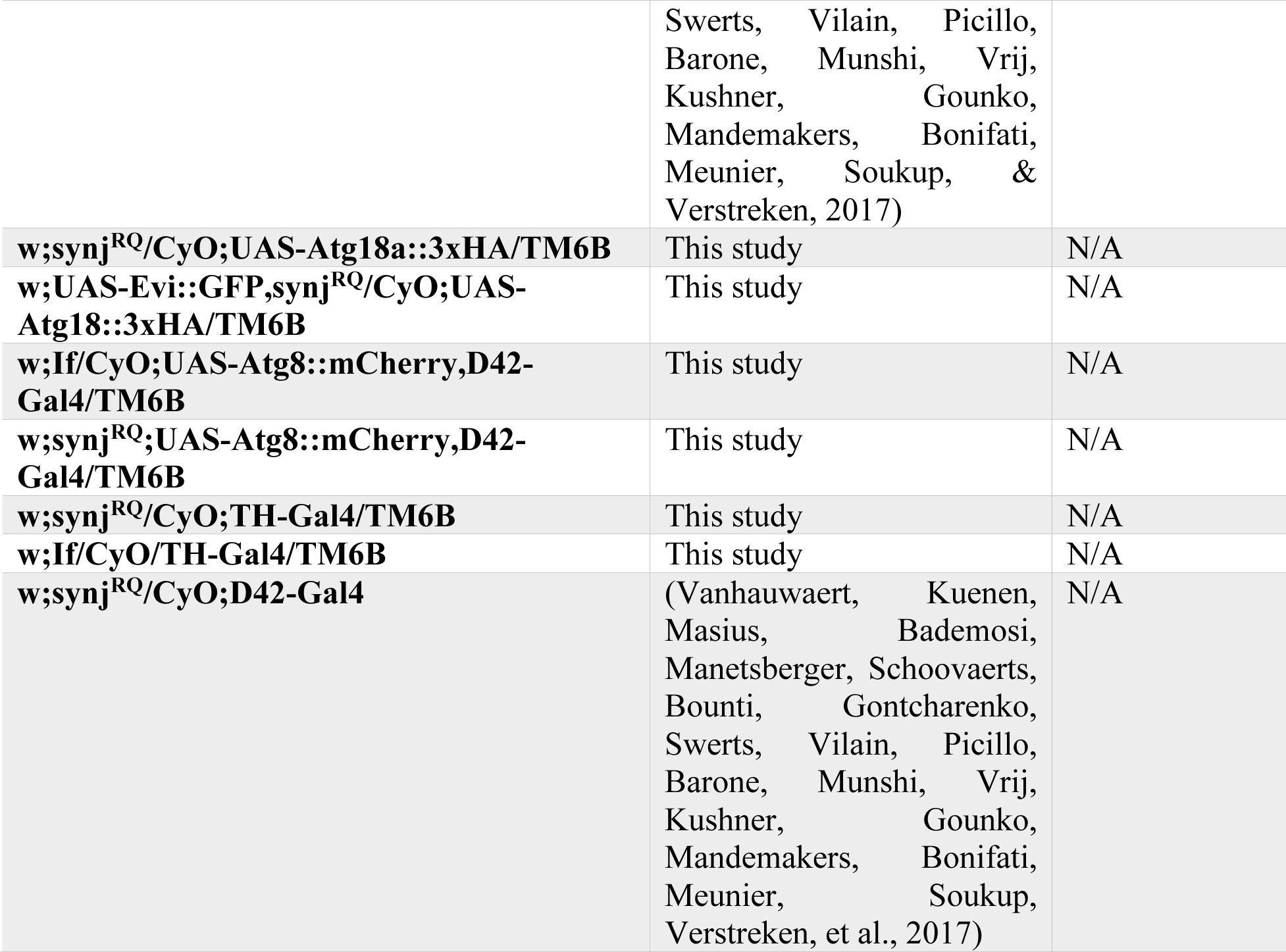

Flies used in this article were grown on standard cornmeal and molasses medium supplemented with yeast at 21 °C. For RNAi mediated knock down experiments flies were raised at 25 °C.

### Antibodies

| Name | Source | Identifier | Dilution |
| --- | --- | --- | --- |
| **Chicken anti-GFP antibody** | AVES Lab | GFP-1020 | 1:1000 |
| **Rabbit polyclonal anti-HRP** | Jackson ImmunoResearch | Cat#323-005-021, RRID:AB_2314648 | 1:500 |
| **Mouse monoclonal anti-DLG antibody** | Developmental Studies Hybridoma (DSHB) Bank | Cat#4F3, RRID:AB_528203 | 1:500 |
| **Rabbit anti-mCherry** | NOVUS | NBP2-25157 | 1:500 |
| **Anti-HA Tag Antibody** | Merck | 05-904 | 1:500 |
| **Rabbit anti-tyrosine hydroxylase (TH)** | NOVUS | NB300-109, lot ajo319o | 1:500 |
| **Mouse antiCD9 APC conjugated (NanoFCM)** | Biolegend | 124812 | 1:1000) |
| **Mouse antiCD63 APC conjugated (NanoFCM)** | Biolegend | 143906 | 1:1000) |
| **Alexa Fluor 488 Goat anti-Chicken antibody Alexa** | Invitrogen | Cat#A-11039, RRID:AB_142924 | 1:1000 |
| **Alexa Fluor 488 Goat anti-mouse antibody** | Invitrogen | A28175 | 1:1000 |
| **Alexa Fluor 555 Goat anti-rabbit antibody** | Invitrogen | Cat#A-21429, RRID:AB_141761 | 1:1000 |

### Larva preparation and immunohistochemistry

Male and female third instar larvae were used indistinctively among the experiments as previously described (Soukup et al., 2016) (Lauwers et al., 2018). The larvae were dissected in fillets using HL3 ((110 mM NaCl, 5 mM KCl, 10 mM NaHCO3, 5 mM Hepes, 30 mM sucrose, 5 mM trehalose, and 10 mM MgCl2, pH 7.2) on sylgard plates. The dissected larvae were fixed with parafolmadehyde 4% in HL3 for 20 minutes at room temperature, washed in PBS (Sigma, P3813-10PAK, Lot #SLCC6084) Phosphate Saline Buffered saline powder, pH 7.4) and then transferred into 0.5 ml tubes. The samples were permeabilised for 1 hour (4 x 15 minutes rotating) with 0.4% triton in PBS (PBST). Then, the samples were blocked using 10% NGS/PBST (blocking solution) for 30 minutes rotating. Samples were incubated with the primary antibody prepared in blocking solution rotating overnight at 4 °C. The next day, samples were washed with PBST for 40 minutes (4 x 10 minutes rotating) and incubated with secondary antibody prepared in blocking solution for 1.5 hour rotating covered in aluminium foil. Then, washing for 1 hour (6 x 10 minutes rotating covered in aluminium foil) and then transferred in PBS. Samples were mounted on Vectashield (Vector Laboratories, ref H-1000, lot ZH0611) and kept at 4°C until imaging.

### Autophagy assay

Starvation assays were performed as previously described (Soukup et al., 2016). Flies were raised at low population density on standard fly food. The day before the experiment around 30 to 40 larvae were randomly selected in order to have low population density grown on standard food supplemental with yeast paste. To induce autophagy, we put the larvae under aminoacid deprivation. For the aminoacid starvation, early third instar larvae were placed on Petri dishes containing 20% sucrose and 1 % agarose during 4 h at 25°C, followed by rapid dissection of the larvae.

For autophagy induction by neuronal stimulation (previously described in (Soukup et al., 2016)), TrpA1 (D42-Gal4)-expressing larvae grown at room temperature 21 °C were placed in a preheated 30 °C tube for 30 min and dissected in heated HL3. Pharmacological autophagy induction via neuronal activity was performed as previously described with minor modifications (Bademosi et al., 2023). In brief, 1 μM Nefiracetam in 0.1% DMSO, 1 mM Ca_2_Cl in HL3 was prepared in 1 ml. For the control, 0.1% DMSO, 1 mM Ca_2_Cl in HL3 was prepared in 1 ml. It was added after the dissection to the sylgard plate with the larvaes for 30 minutes. After that fixation in PFA was done and continued the protocol as usual. For the chloroquine assay, early third-instar larvae were fed for 4 h at 25°C on standard food containing 10 mg/ml chloroquine disphosphate salt (Sigma, C6628-25G)

### Image acquisition and quantitative image analysis

For imaging, confocal microscopy (TCS SP5, Leica Microsystems) with a Leica HCX PL APO lambda blue 63×/1.40-0.60 OIL UV objective was used. In particular, the settings used to obtain the data were: image size 1024×1024 pixels 12 bits. The pinhole size was fixed at 1 AU. Samples from 5 larvae per genotype and condition were imaged using a single confocal plane 4-6 neuromucular junction (NMJ) per larvae. In order to guarantee representability, each NMJ image was taken from different segment (A2, A3 or A4) using muscle 12 or 13 and 2 or 3 from each side (left-right) of the larvae. Larvaes were scanned under the same settings. Quantification of intensity or number of puncta was done in NIH ImageJ. Exosomes and dots were quantified using the previously published automatic method exoquant or autophagoquant (Sanchez-Mirasierra et al., 2021)

Colocalization of Evi::GFP or Evi::Cherry positive dots with Autophagic makers (Atg8::mCherry, Atg3::HA, Atg5::HA, Atg16::GFP, Atg17::GFP and Atg18::GFP) was calculated using JaCoP in imageJ, where an object-based Pearson correlation coefficient is defined by thresholding the puncta (thresholded overlay colocalization coefficient).

All images were processed using ImageJ and figures were composed using Adobe Illustrator 2021.

### Exosomes extraction from fly brains

We isolated exosomes from adult fly brains through multiple ultracentrifugation steps followed by a sucrose density ultracentrifugation allowing the fractionation of extracellular vesicles depending of their density. In detail, adult flies with the following genotypes (w^1118^; C155/+;UAS-EVI::GFP/+ and C155/+;UAS-EVI::GFP,synj^RQ^/ synj^RQ^) were frozen in liquid nitrogen and stored at -80 °C until the day of exosome extraction. 220 flies/genotype were introduced into liquid nitrogen followed by immediately vortexing at maximum speed for decapitation. Heads were separated from the body by sieving through Nitex-Nylon meshes with different size (700um, 630um and 15um). The heads were transferred with a funnel into a 5ml Eppendorf and weighted (around 0.06 ± 0.01 g). Then, 500 μl of HL3 was added and samples were homogenized on ice using a motorized pestle during 3 minutes. The homogenate was incubated with collagenase type 3 (Worthington, #LS004182, lot 42B22080) 1.64U /mg of head (1.64 U/μl) for 20 minutes at 37°C (for example: 1.5 mg of collagenase (405U) were diluted in 250 μl HL3, and then 1 μl of collagenase/HL3 solution per 1 mg of head). During the 20 minutes incubation, after 5 minutes the samples were vortexed, then after 10 more minutes (15 from the total) the samples were resuspended by pipetting with the tip of the pipette cut off. The reaction was stopped using Protease inhibitor (Fisher Scientific, ref. 10444414, lot VH307175 Halt^TM^ Protease Inhibitor Single use Cocktail (100x)) diluted on ice cold HL3 to obtain 2x 500 μl (inhibitor/HL3) that were added to the other 500 μl (heads/HL3). The samples were passed through a cell strainer 70 μm into a 50 ml falcon tube. The samples were centrifuged at 300 x g for 10 minutes at 4 °C. The pellet contains the nuclei. Supernatant was transferred to a new 15 ml falcon tube and centrifuged for 2000 g for 10 minutes at 4 °C. The pellet contains microsomes. The supernatant was then transferred to an ultracentrifuge tube.

EBSS (Fisher Scientific, ref. 11570466, (1X)) was added to the tube to complete the tube (around 4ml). Then, the tubes were weighted in order to have the same weight (± 0.1 g). Additional EBSS was used to balance the tubes when necessary. The samples were ultracentrifuged at 100 000 g for 70 minutes at 4 °C. The supernatant was discarded. The pellet was resuspended in 2 ml of 0.475M of sucrose solution and give a quick vortex. 2M sucrose solution in PBS was prepared (171.2 g sucrose in 250ml PBS) and from that sucrose solution other sucrose solutions: 1.5 M, 1 M, 0.825 M, 0.65 M and 0.475 M were made. Then, a sucrose step gradient six-2ml steps starting from 2M to 1.M, 0.825 M, 0.65 M and 0.475 M was made. Lastly the 0.425 M fraction containing the resuspended pellet was added on top, you can use the 0.425 M sucrose solution to calibrate for the ultracentrifuge. The gradient tubes were centrifuged at 200 000 g for 20 hours at 4 °C

The fraction III (1ml) was removed, the next fractions are 2ml in volume except the last one that is 1 ml also. Fractions III contains the lighter particles and fraction IX the denser. Each fraction was diluted in 10 ml of PBS, the adjustment for them to weight the same were made with PBS to have an error of ± 0.1 g. Ultracentrifugation was at 100 000 g at 4 °C for 70 minutes for each fraction. The supernatant was discarded and the pellet was resuspended in 40 μl of cold PBS. The fractions were stored at -80 °C.

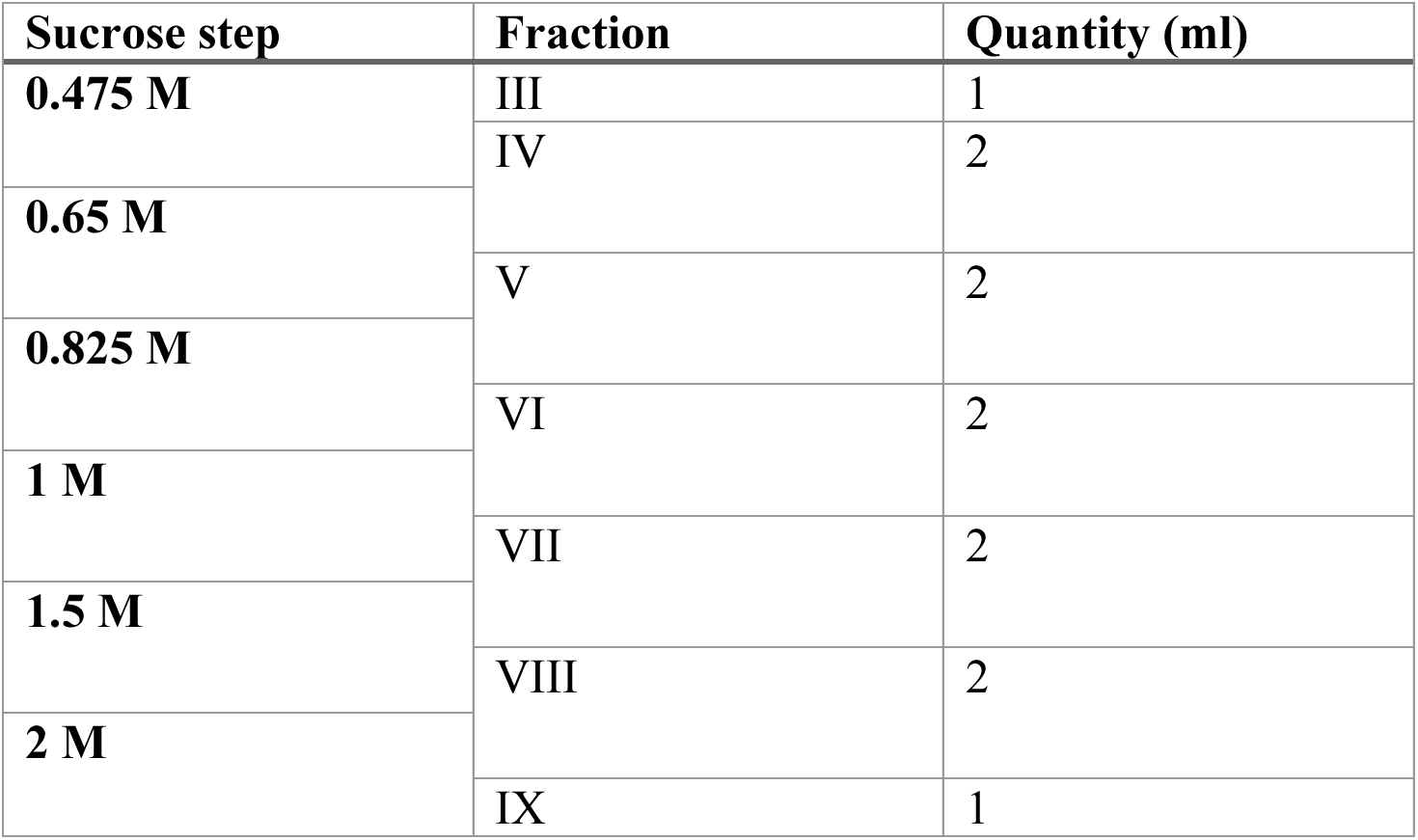

### NanoFCM flow cytometry

The size, concentration and GFP specificity of EV samples was characterized using the flow NanoAnalyzer (NanoFCM Co.). Briefly, NanoFCM quality control nanospheres (NanoFCM Co. Ltd, ref. QS2503) with a size of 250 nm and concentration of 1.99E+10 particles/mL were diluted 1:100 in ultrapure Milli-Q water for a final volume of 100 µl in order to align and calibrate the NanoAnalyzer’s 488 nm and 640 nm lasers and SPCM detectors. The diluted nanospheres were analyzed for 1 minute at a sampling pressure of either 1.0 kPa or 1.5 kPa. Following calibration, NanoFCM silica nanospheres cocktail (NanoFCM Co. Ltd, ref S16M-Exo) containing a mixture of monodisperse silica nanospheres with diameters (68, 91, 130 and 155 nm) were used for extrapolation of size information of EV samples based on the side scatter intensity. The diluted nanospheres were analyzed for 1 minute at a sampling pressure of either 1.0 kPa or 1.5 kPa. For acquisition of size, concentration and GFP specificity of EVs, samples were diluted between 1:5 and up to 1:200 in 1X PBS (filtered with a 0.22 µm filter) and analyzed for 1 minute at a sampling pressure of 1.0 kPa. For phenotyping of EVs, 4 µl of EV sample was incubated for 30 min with 1 µl of anti-CD9 APC or anti-CD63 APC in the dark at room temperature. Stained samples were further diluted to 1:1000 with 1X PBS (filtered with a 0.22 µm filter) and analyzed with the 640 nm laser for 1 minute at a sampling pressure of 1.5 kPa. Unstained and PBS-antibody controls were used to determine specificity of staining.

### Transmission Electron Microscopy

An aliquot of EV samples (5 µl) were fixed with 4% paraformaldehyde (filtered with 0.22 µm filter) on ice. The fixed sample (10 μl) was be absorbed onto glow-discharged 300-mesh heavy duty carbon-coated formvar Cu grids (ProSciTech, Kirwan, QLD) for 20 min followed by washing with PBS (3 x 5 min). The grids were further incubated with 1% glutaraldehyde for 5 min and washed with Milli-Q water (8 x 2 min). Subsequent negative staining was carried out by incubating the grids with uranyl-oxalate (pH 7) in the dark for 5 mins followed by incubation with methyl cellulose-uranyl acid solution in the dark on ice for 10 mins. High-contrast images were acquired using a HITACHI H7650 transmission electron microscope (Hitachi, Krefeld, Germany). The electron microscopy was performed at the Bordeaux Imaging Centre.

### Behavioral tests

Adult climbing ability (also known as negative geotaxis climbing ability) was tested accordingly to (Wilson et al., 2020) (Nichols et al., 2012). In brief, newly emerged Drosophila adults were selected and aged at 25°C with a 16h/8h day/night cycle. The day before the experiment, 25 adult flies (12 male and 13 female) of the same age and genotype were collected and transfer into a new tube containing fresh food without using CO_2_. Before the experiment I took flies from the incubator and allowed flies to acclimate to the environment (22°C and 45% HR) for 15 min. Flies were then placed into an empty plastic vial (25mm diameter) where I previously register the mark of 6cm from the bottom with a line. Flies were gently tapped to the bottom of the vial and we quantified the number of flies that crossed the line within 10s (+1). Flies that do not cross the line count 0 and flies that cross back or fall after crossing the line count -1. A minimum of 6 consecutive trials were performed, each followed by a rest time of 1 minute and using fresh vials and percentage of flies crossing the 6cm mark was calculated. Before each experiment the group of flies were inspected using the scope and death flies or adults showing body defects that could prevent normal climbing ability (broken legs or wings) were discarded. To avoid cofounds, experiments were performed at the same place to guarantee similar light, temperature and humidity conditions and around the same time of the day to maintain uniform circadian rhythms.

The Bang-sensitivity behavioral test measuring recovery from seizure-like behaviors upon mechanical stimulation was performed using a modified version of the protocol described in Tao et al (Tao et al., 2011). In brief, newly emerged flies were selected and aged at 25°C with a 16h/8h day/night cycle. The day before the experiment, 20 to 25 adult flies (12 male and 13 female) of the same age and genotype were collected and transfer into a new tube containing fresh food without using CO_2_. Before the experiment, the experimental flies were acclimated to the environment (22°C and 45% HR) for 15 min. Flies were then placed into a clean plastic vial (25mm diameter) where I previously register the mark of 6cm from the bottom with a line. Vials containing the adult flies were mechanically stimulated by vortexing at maximum power (VWR, Ref. 444-1372. 51W) for 10 seconds (we previously tested that mechanical stimulation by vortexing up to 20 seconds has no effect in lethality in our genotypes). After vortexing, vials were placed on a flat bench and recorded for 1 minute. Videos were then analyzed and two parameters were quantified. First, the climbing ability to cross a mark placed at 6cm from the bottom in 10 seconds. Second, I measured their recovery “off bottom” if they were recovered from seizure paralysis and started coordinated movement behavior quantified as their ability to leave the bottom of the vial at defined time points during 60 secs.

### Dopaminergic neurodegeneration assay

1–2-day old flies were aged for 10 days at 25°C and transferred every 3 days to fresh media. At day 10, fly brains were dissected in the physiological solution HL3, fixed for 30 min. in 4% PFA. Subsequently brains were washed in PBS and incubated in 10% NGS-PBT (0.4% Triton in PBS) for 1h and stained with mouse anti- tyrosine hydroxylase (TH) antibody (1:500, Millipore) in 10% NGS-PBT overnight at 4°C. Before imaging samples were mounted in Vectashield (Vector Laboratories). Images of dopaminergic clusters were captured with confocal microscopy (TCS SP5, Leica Microsystems) with a 20×/0.70 objective. The total number of TH-positive neurons (dopaminergic) present in both the hemispheres for PPM3 and PPL1 clusters were counted manually.

### Statistics

Distribution of data was analyzed using a D’Agostino-Pearson Omnibus test. Normal data were tested with parametric tests: for two datasets we used a student’s t test. For more than two datasets, we used a one-way analysis of variance test (ANOVA) and a post hoc Tukey test. Non-normal distributed data were tested using non-parametric tests: for two datasets we used a Mann Whitney test and for more than two datasets an ANOVA-Kruskal Wallis test and a Dunn’s test was used. Significance of statistical difference was defined as **** = p<0.0001, *** =p<0.001, **=p<0.01, *=p<0.05, ns= p>0.05. The number of analyzed samples (animals, NMJ boutons or brains) are indicated in the figure legend as N (no. of animals) and n (no. of analyzed images).

### Software

a. ImageJ/FIJI (https://imagej.net/Fiji) (Schindelin et al., 2012)
b. Exoquant and Autophagoquant detector macros for ImageJ (Sanchez-Mirasierra et al., 2021)
c. Excel (Microsoft https://www.microsoft.com/es-ww/microsoft-365/excel), Prism 9 (Graphpad https://www.graphpad.com/scientific-software/prism/) and Illustrator 2021 (Adobe, https://www.adobe.com/es/products/illustrator) Alternatively, you can use R (The R Foundation, https://www.r-project.org/)
d. NF Profession 1.0
e. Digitalmicrograph software (https://www.gatan.com/products/tem-analysis/gatan-microscopy-suite-software)

## AUTHOR CONTRIBUTIONS

SFS conceived the study, IS-M, SFS, SH-D wrote the paper, IS-M, SFS, SH-D analyzed data. IS-M SH-D, SG, LB-C, J-C D, AAM and SFS, performed experiments and/or provided reagents.

## ACKNOWLEDGMENTS

We thank V. Budnik, M. Boutros, T. Neufeld, G. Juhasz, the Bloomington Drosophila Stock Center, FlyORF and the Developmental Studies Hybridoma bank for reagents.

SFS received financial support from the PID2023-153022NB-100 project financed by MCIN/AEI/10.13039/501100011033 /FEDER, EU, Ikerbasque Foundation for Science and the Region Nouvelle-Aquitaine. SFS acknowledge support from Marie Sklodowska-Curie COFUND Programme of the European Commission HORIZON-MSCA-2022-COFUND-101126600-SmartBRAIN3. JCD received support from the Région Nouvelle-Aquitaine CHESS Exomarquage 13059720-13062120, and the ANR-23-CE14-0056-01.

